# Epithelia are multiscale active liquid crystals

**DOI:** 10.1101/2022.02.01.478692

**Authors:** Josep-Maria Armengol-Collado, Livio Nicola Carenza, Julia Eckert, Dimitrios Krommydas, Luca Giomi

## Abstract

Biological processes such as embryogenesis, wound healing and cancer progression, crucially rely on the ability of epithelial cells to coordinate their mechanical activity over length scales order of magnitudes larger than the typical cellular size. While regulated by signalling pathways, such as YAP (yes-associated protein), MAPK (mitogen-activated protein kinase) and Wnt, this behavior is believed to additionally hinge on a minimal toolkit of physical mechanisms, of which liquid crystal order is the most promising candidat. Yet, experimental and theoretical studies have given so far inconsistent results in this respect: whereas *nematic* order is often invoked in the interpretation of experimental data, computational models have instead suggested that *hexatic* order could in fact emerge in the biologically relevant region of parameter space. In this article we resolve this dilemma. Using a combination of *in vitro* experiments on Madin-Darby canine kidney cells (MDCK), numerical simulations and analytical work, we demonstrate that both nematic and hexatic order is in fact present in epithelial layers, with the former being dominant at large length scales and the latter at small length scales. In MDCK GII cells on uncoated glass, these different types of liquid crystal order crossover at 34 ***µ***m, corresponding approximatively to clusters of 21 cells. Our work sheds light on the emergent organization of living matter, provides a new set of tools for analyzing the structure of epithelia and paves the way toward a comprehensive and predictive mesoscopic theory of tissues.

---

Detecting orientational order in epithelia [1, 2] has been the focus of several recent studies [3–6]. The task is commonly approached by tracking the longitudinal direction of individual cells by diagonalizing a rank−2 tensor − i.e. the so called structure tensor [7] or equivalently the shape tensor [8, 9] in case of segmented images − that embodies the geometry of the polygonal cells (Fig. 1a). The resulting two-dimensional orientation field is then used to identify topological defects [3–6], which in turn provide a fingerprint of the underlying orientational order. Liquid crystal defects (also known as disclinations) are isolated singularities in the orientational field and can be classified according to their winding number or “strength” *s*, defined as the number of revolutions of the orientation field along an arbitrary contour encircling the defect core [10]. Because in a two-dimensional liquid crystal with *p*−fold rotational symmetry (i.e. symmetry under rotations by 2*π/p*) this number must be an integer multiple of 1*/p*, defects such as vortices, asters and spirals, for which *s* = 1, are a signature of a polar phase (i.e. *p* = 1); comet- and star-shaped disclinations, whose winding numbers are *s* = 1*/*2 and *s* = −1*/*2 respectively, are representative of a nematic phase (i.e. *p* = 2); whereas 5–fold and 7 fold disclinations, with *s* = 1*/*6 and *s* = −1*/*6, are the elementary topological defects in hexatics (i.e. *p* = 6).

**Figure 1.**
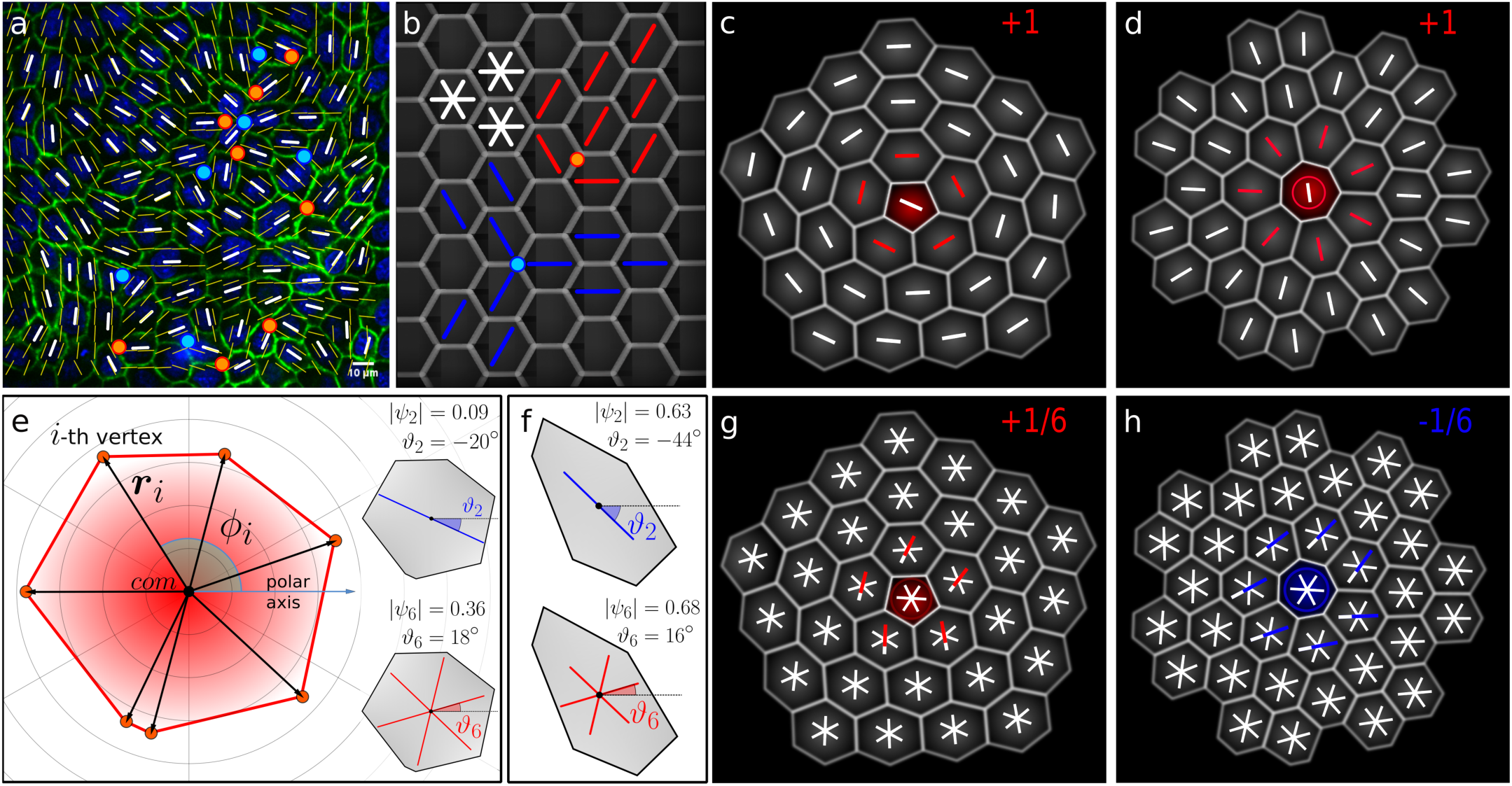
Topological defects and *p−*fold rotational symmetry. **a** A typical configuration of the nematic orientation field (white rods), obtained from a sample of MDCK GII cells upon diagonalizing the shape tensor [8, 9]. Yellow rods represent the interpolated nematic field. Here and in the following positive and negative defects are marked in red and blue respectively, regardless of the magnitude of their winding number. **b** Because of the 6−fold symmetry of regular hexagons, there is no well defined longitudinal direction, thus it is possible to construct a defective configuration, featuring a pair of ± 1*/*2 disclinations, even though the lattice is defect free. **c**,**d** Disclinations in a hexatics consists of pentagonal (i.e. *s* = 1*/*6) and heptagonal (i.e. *s* =−1*/*6) site embedded in an otherwise 6−fold background. Attempting to detect these elementary defects by tracking the longitudinal direction of the cells (with rods), correctly yields a defect at the center of the clusters, however, because of the mismatch between the 6 fold symmetry of the configuration and the 2*−*fold symmetry of the order parameter both defects are detected with the incorrect winding number *s* = 1. **e** Graphical representaiton of the *p−*fold order parameter, Eq. (2), for a generic polygon (heptagon). The quantities ***r***_*i*_ = {*x*_*i*_, *y*_*i*_} and *ϕ*_*i*_ = arctan(*y*_*i*_*/x*_*i*_) represent, respectively, the position of the vertices of the polygon with respect to its center of mass (i.e. *com*) and their orientation with respect to the horizontal direction (i.e. polar axis). Inset shows the nematic (top) and hexatic (bottom) order parameter *ψ*_*p*_ superimposed on the polygonal shape of the main panel. **f** Example of the *ψ*_*p*_ order parameter, Eq. (2), for an elongated hexagon. The irregular heptagon in panel (e) is closer in shape to a regular hexagon, thus the order parameter *ψ*_6_ is an order of magnitude larger than *ψ*_2_. The outcome is reversed in the irregular hexagon in panel (f), which, as a consequence of its elongation and despite being 6–sided, yields *ψ*_2_ *> ψ*_6_. In both panels, the blue rods and the 6−legged stars corresponds respectively to the 2*−*fold and 6−fold orientations of the polygons and are oriented in such a way that maximizes the overall probability of finding a vertex in the direction of the legs. **h**,**i** Correct recognition of the hexatic disclinations shown in panels (c) and (d) using *ψ*_6_. In both panels one of the legs of the order parameter has been colored as a guide to the eye. By following the order parameter along a positive oriented (anticlockwise) close loop encircling the defect core, the red leg rotates anticlockwise for the positive defect in panel (e). After an full rotation, the colored legs rotates of an angle 2*π/*6 corresponding to a winding number *s* = 1*/*6. Conversely, in panel (i) the blue leg rotates clockwise and covers and angular displacement of *−π/*3 corresponding to a winding number *s* = *−*1*/*6.

Although inferring order from defects represents a consolidated strategy in liquid crystals science since the times of Georges Friedel [11] − who used it to decipher and classify phases such as nematic, cholesteric, and smectic − this specific protocol, based on tracking the cells’ longitudinal direction, becomes progressively less reliable as *p* increases. To illustrate this issue we show in Fig. 1b how applying the same protocol to a perfect honeycomb lattice can lead to the misdetection of a pair of ± 1*/*2 nematic disclinations. This originates from the fact that, while regular hexagons are invariant under rotations by 60^°^, the orientation field constructed from the longitudinal direction of hexagonal cells cannot discriminate between the three equivalent directions defined by pairs of opposite vertices. Similarly, in Figs. 1c and 1d we show how detecting an elementary hexatic diclination correctly yields a topological defect, but with incorrect winding number *s* = 1.

To overcome this difficulty, here we introduce a generalized rank−*p* shape tensor, able to capture arbitrary *p*−fold rotational symmetries, with *p* any natural number. Given the polygonal contour of a cell, whose *V* vertices have coordinates ***r***_*i*_ = {*x*_*i*_, *y*_*i*_} with respect to the cell’s center of mass (Fig. 1e), our generalized shape tensor can be defined as

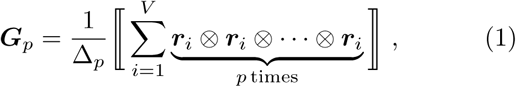

Where 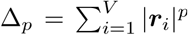, and the operator ⟦…⟧ has the effect of rendering its argument symmetric and traceless [12]. For *p* = 2, Eq. (1) gives, up to a normalization constant, the traceless part of the standard rank−2 shape tensor [8, 9]. Regardless of its rank, the tensor ***G***_*p*_ has only two linearly independent components in two dimensions [13, 14], from which one can extract information about the cells’ orientation and anisotropy. In particular, using a generalization of the spectral theorem to tensors with arbitrary rank [15, 16], one can show that all elements of ***G***_*p*_ are proportional to either the real or the imaginary part of the complex order parameter

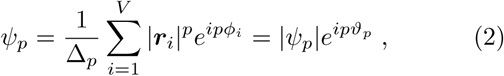

where *ϕ*_*i*_ = arctan(*y*_*i*_*/x*_*i*_) the angular coordinate of the *i−* th vertex of a given cell (Fig. 1e). The angle *ϑ*_*p*_, on the other hand, corresponds to the *p−*fold orientation of the whole cell with respect to the horizontal direction. In practice, this is equivalent to the inclination of a *p*–legged star centred at the cell’s center of mass and oriented in such a way to maximize the probability of finding a vertex in the direction of either one of the legs. Some example of this construction is shown in Fig. 1e and 1f, where *ψ*_*p*_ is computed for more or less elongated irregular polygons. When applied to defective configurations, our method yields the correct winding numbers *s* = 1*/*6 (Fig. 1g and 1h).

With the tensor ***G***_*p*_ in hand, we next investigate the emergent orientational order in confluent monolayers of MDCK GII cells (Figs. 2a and 2b). After segmenting the images, by taking advantage of the previous labeling of E-cadherin, we track the cells’ contour and from the coordinates of the vertices we compute the order parameter *ψ*_*p*_, Eq. (2). We analyze a total of 68 images of confluent monolayers (see the Methods for details) with each one of them comprising 140 ± 31 cells (mean ± s.d.). Fig. 2c shows the probability distribution of *ψ*_*p*_ for *p* = 2 and 6. Interestingly, the distribution of *ψ*_6_ is symmetric and spreads over a broad range of values; conversely the distribution of *ψ*_2_ features a peak at approximatively 0.35, with a decreasing tail at larger values. The MDCK GII cells analyzed in this study are, therefore, more prone to arrange in isotropic rather than elongated shapes. This results in a disordered and yet orientationally coherent tiling of the plane, where a majority of hexagons coexists with large minorities of pentagons and heptagons, as indicated by the distribution of the number of neighbors in Fig. 2d. We compare these observations with numerical simulations of two different theoretical models of epithelia: i.e. a continuous multiphase field model (mpf) [17–19] (Fig. 2e) and the discrete Voronoi model [20–23] (Fig. 2f), both in qualitative agreement with experimental data.

**Figure 2.**
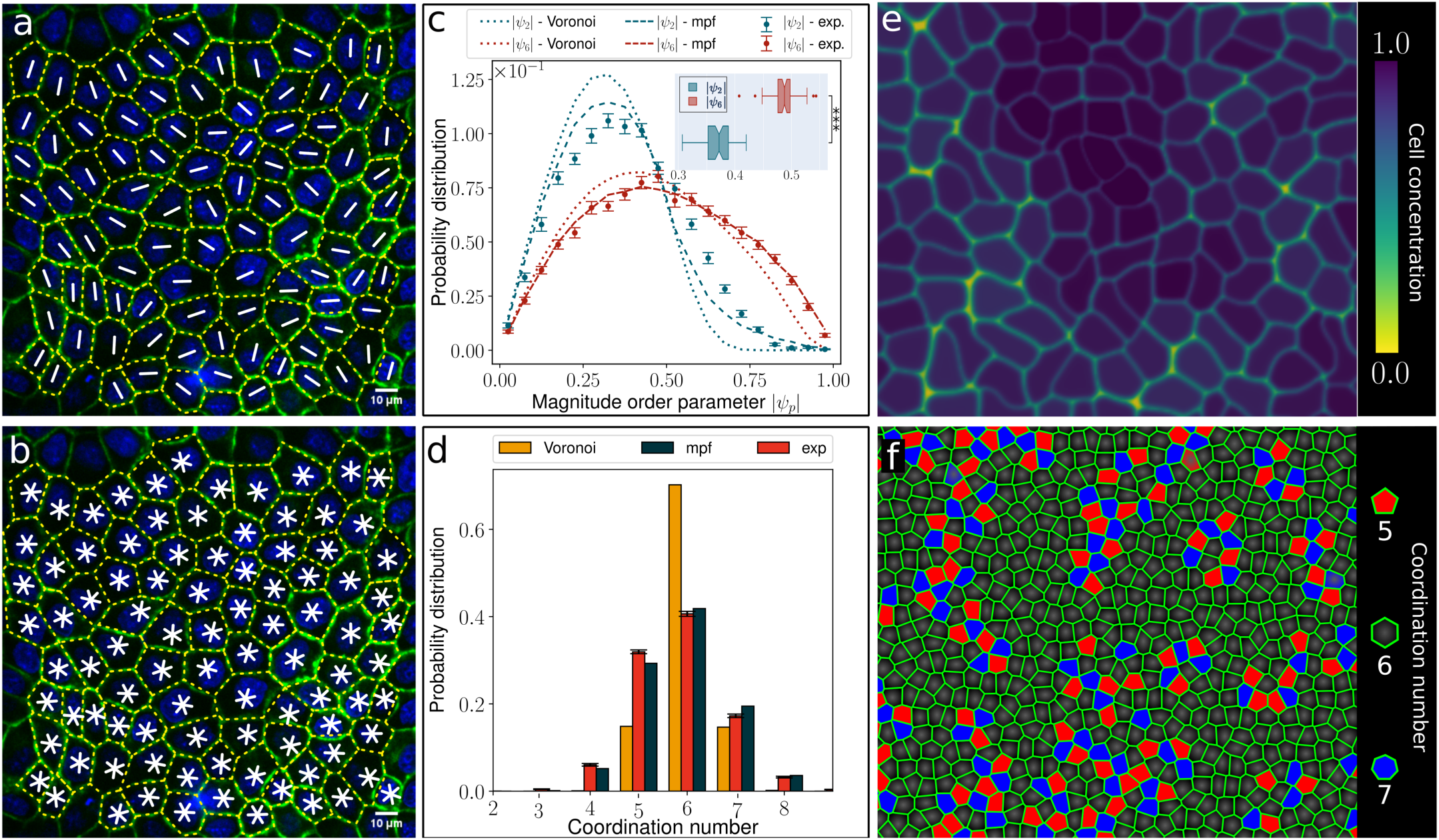
Symmetry of MDCK cells in confluent monolayers. **a**,**b** Confocal image of a confluent MDCK GII monolayer (green, E-cadherin and blue, nuclei). The dashed yellow lines trace the contour of the cells as identified after image segmentation. The white rods **a** and stars **b** respectively mark the 2*−*fold and 6*−*fold orientation of cells and have been obtained from the order parameter *ψ*_*p*_, Eq. (2). **c** Probability distribution of the magnitude of the order parameter |*ψ*_*p*_| for *p* = 2 (blue) and *p* = 6 (red). Experimental data points are obtained by averaging over 68 different images with each containing 140±31 (mean ± s.d.) cells. The mean value of the distributions are ⟨*ψ*_2_ ⟩ = 0.370± 0.030 (mean ± s.d.) and ⟨*ψ*_6_ ⟩ = 0.49 ±0.05 (mean ± s.d.). The boxplot in the inset shows the average magnitudes of the order parameters of 68 imaged monolayers. ⟨ *ψ*_2_ ⟩ and ⟨ *ψ*_6_ ⟩ are significantly different with a p–value of p *<* 10^*−*4^, calculated by using the two-sided Wilcoxon rank sum test. Dashed and dotted lines are obtained from numerical simulations of the multiphase-field (mpf) and Voronoi models. **d** Probability distribution of cell coordination number for experiments and simulations. The hight of the bar represents the mean of 68 analyzed images. The mean values of the coordination number distributions are 5.8 ± 0.9 (mean ± s.d.) for experiments and 5.9 ± 0.9 (mean ± s.d.) for multiphase field simulations and 6.0 ± 0.6 (mean ± s.d.) for Voronoi simulations. In **c** and **d**, error bars are computed from the standard error of mean. **e** Contour plot of the local cell concentration of a multiphase-field simulation with 360 cells in a magnified region of the simulation box showing approximately one third of the system. Darker regions correspond to areas dense with cells and lighter regions to areas where cells are sparser (see legend box). **f** Configuration of a numerical simulation of the Voronoi model. The cells in red (blue) have 5 (7) neighbors while others have 6.

In order to quantify the amount of orientational order in the system, we next compare the orientation of each cell with that of its neighbors, by means of the following coarse-graining procedure. Given a disk Ω_*R*_ = Ω_*R*_(***r***), with radius *R* and centred at ***r***, and letting ***r***_*c*_ be the position of the center of mass of the *c −* th cell, we define the coarse grained order field Ψ_*p*_ = Ψ_*p*_(***r***) as

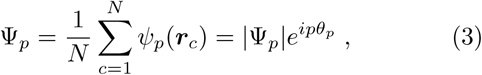

where *N* is the number of cells whose centre of mass lies within Ω_*R*_, while |Ψ_*p*_| = |Ψ_*p*_(***r***)| and *θ*_*p*_ = *θ*_*p*_(***r***) are respectively the magnitude and phase of the complex order parameter Ψ_*p*_, conveying information about the amount and direction of *p*–fold orientational order at the length scale *R* (Fig. 3a).

**Figure 3.**
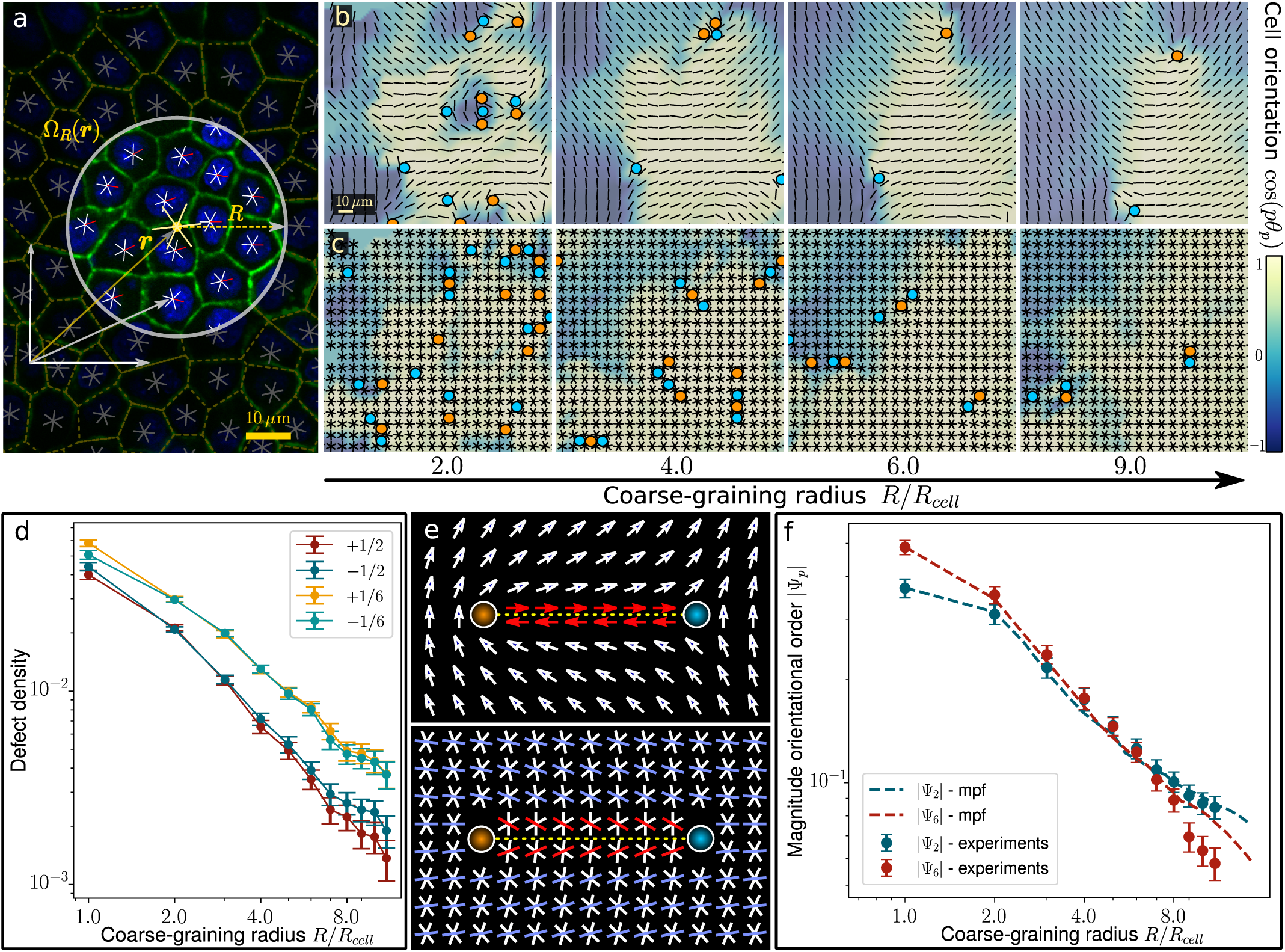
Coarse-graining and multiscale features of confluent cell monolayers. **a** Illustration of the coarse-graining procedure entailed in Eq. (3). A disk Ω_*R*_ = Ω_*R*_(***r***) (encircled in gray), with radius *R* and centered at the point ***r*** (in general not coincident with the center of mass of any specific cell) is superimposed to a segmented image of the cell monolayer and the cells in its interior are used to compute the coarse grained filed Ψ_*p*_. The large yellow star at the centre of the disk shows the orientation of the whole cluster. **b, c** Nematic (top row) and hexatic (bottom row) coarse grained fields Ψ_2_ and Ψ_6_ versus the coarse graining radius *R*, expressed in units of the average cell size *R*_cell_ = 7.4 *µ*m. In both panels positive and negative defects are marked in red and blue respectively (± 1*/*2 for nematic and ± 1*/*6 for hexatic). **d** Defect density at varying the coarse graining radius *R*. **e** A mismatch between the defect charge and the symmetry of the *p*–atic liquid crystal gives rise to unphysical singular line (see Sec. SII in Ref. [16]). Top (bottom) panel shows a pair of nematic (hexatic) defects of winding number *s* = ±1*/*2 (*s* = ±1*/*6) **f** Magnitude of Ψ_2_ and Ψ_6_ versus the coarse graining radius *R* measured from experimental and numerical mpf data. Both data sets fit the power law 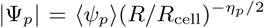, with *η*_*p*_ a non-universal exponent [13, 14], with the following fitting parameters: (experiments) *η*_2_ = 0.41 ± 0.01, *η*_6_ = 0.49 ± 0.01; (mpf) *η*_2_ = 0.43 ± 0.02, *η*_6_ = 0.48 ± 0.01. In both experiments and multiphase field simulations, the |Ψ_2_| and |Ψ_6_| order parameters crossover at the length scale *R*_*×*_, with: (experiment) *R*_*×*_*/R*_cell_ = 4.6 ± 1.0; (mpf) *R*_*×*_*/R*_cell_ = 5.0 ± 1.2. In **d** and **f** the error bars correspond to the standard error on the mean.

The outcome of this analysis is shown in Figs. 3b and 3c. At length scales comparable with the average cell size − i.e. *R ≲ R*_cell_, with *R*_cell_ = 7.4 ± 1.9 *µ*m the average cell radius computed as half of the distance between the cells’ centers of mass−both the nematic (Fig. 3b) and the hexatic (Fig. 3c) coarse grained fields are populated by topological defects. For *p* = 6, in particular, the monolayer appears organized into regions characterized by spatially uniform hexatic order, separated by arrays of ±1*/*6 disclinations, similarly to grains and grain boundaries in polycrystals [24]. Increasing *R* has the effect of smoothing the Ψ_6_ field, thereby absorbing neutral pairs of disclinations into a gently varying 6−fold orientation field, resulting in a power law decreasing defect density (Fig. 3d).

The scenario differs dramatically for *p* = 2 (Fig. 3b). In this case, many of the defective structures identified in the configuration of the hexatic field at the small length scales are replaced by very sharp and yet defect-free textures. This peculiarity originates precisely from the mismatch between the actual 6*−*fold symmetry of the configuration at the cellular scale and the 2*−*fold symmetry of the order parameter used to describe it, in a similar fashion as using a (polar) vector field to describe a nematic disclination gives rise to singular lines where the polar field “jumps” by an angle *π* (Fig. 3e). Conversely, at larger length scales, the majority of nematic defects is replaced by regions where the nematic field Ψ_2_ smoothly varies across the sample, with exception for a small number of isolated ±1*/*2 disclinations (Fig. 3b and 3d). These observations are further supported by the scaling behavior of the magnitude of the fields Ψ_2_ and Ψ_6_ as the coarse-graining radius *R* varies (Fig. 3f). In particular, both | Ψ_2_| and | Ψ_6_ | are finite at all length scales in the range 1≤ *R/R*_cell_ *<* 10, but, while | Ψ_6_| is prominent at small length scales, this is overweighted by | Ψ_2_| at large length scales. For our MDCK GII cells on uncoated glass, the crossover occurs at *R*_×_*/R*_cell_ = 4.6 ±1.0, corresponding to clusters of approximatively 21 cells. The same crossover is also observed in our numerical simulations of the multiphase field model, with the crossover scale *R*_×_*/R*_cell_ = 5.0± 1.2, while it is not found in simulations of the Voronoi model, where hexatic order is dominant at all length scales (Fig. 3f).

Taken together, our experimental and numerical results demonstrate that epithelial monolayers behave as multiscale active liquid crystals, with 6–fold hexatic order characterizing the spatial organization of the cells at small length scales, while nematic order dictates the large scale structure of the monolayer. The crossover length scale is, as intuitive, non-universal, but depends on the molecular repertoire and the material properties of the specific phenotype, as well as on the mechanical properties and the surface chemistry of the substrate.

In conclusion, we have investigated the existence of orientational order in epithelial layers, as a possible route toward complementing the complex network of regulatory pathways that tissues have at their disposal, with a minimal toolkit of physical mechanisms, whose primary effect is to coordinate the activity of individual cells to achieve multicellular organization. Upon introducing a novel tensorial descriptor of cellular orientation *−*i.e. the generalized shape tensor ***G***_*p*_ − we have demonstrated that multiple types of liquid crystal order can coexist in epithelial layers at different length scales. In particular, hexatic order (i.e. *p* = 6) is prominent at the small scale (i.e. in clusters of up to 21 cells in our MDCK GII samples), whereas nematic order (i.e. *p* = 2) characterizes the structure of the monolayer at larger length scales. This novel approach creates the basis for a correct identification of topological defects − whose biophysical role in epithelia has recently focused great attention [3–5], especially in the context of morphogenesis [25–28] − and further provides the necessary knowledge for the foundation of a comprehensive and predictive mesoscopic theory of collective cell migration [29]. In addition to advancing current techniques for the interpretation of *in vitro* experimental data, our findings highlight a number of potentially crucial properties of epithelia *in vivo*. First, collective cell migration in epithelia relies on both remodelling events at the small scale − such as cell intercalation and the rearrangement of multicellular rosettes [30, 31] − as well as large scale flows [26]. Therefore the underlying *hexanematic* multiscale organization and the specific magnitude of the crossover scale *R*_×_ are expected to have a profound impact on how the geometry of the environment affects the specific migration strategy. E.g. metastatic cells traveling through micron-sized channels in the extracellular matrix during cancer invasion [32] will more likely rely on local hexatic-controlled remodelling events, whereas unconfined wound healing processes [33] are more likely to leverage on system-wide nematic-driven collective flows. Second, as both hexatic and nematic liquid crystals can feature topological defects, these are expected to interact, thereby affecting processes such as the extrusion of apoptotic cells [4], the development of sharp features during morphogenesis [28, 34] and, in general, any remodelling or morphogenetic event that can take advantage of the persistent pressure variations introduced by active defects [5]. Finally, in the light of what said above, it is evident that understanding how the crossover scale *R*_×_ can be controlled, either chemically or mechanically, may ultimately represents the key toward deciphering tissues’ collective dynamics.

## ACKNOWLEDGEMENTS

This work is supported by the European Union via the ERC-CoGgrant HexaTissue (L.N.C., D.K. and L.G.) and by Netherlands Organization for Scientific Research (NWO/OCW) as part of the research program “The active matter physics of collective metastasis” with project number Science-XL 2019.022 (J.-M.A.-C and L.G.). Part of this work was carried out on the Dutch national e-infrastructure with the support of SURF through the Grant 2021.028 for computational time. J.E. and L.G. acknowledge M. Gloerich, UMC Utrecht, for providing us the MDCK cells. All authors acknowledge Ludwig Hoffmann for fruitful discussions.

## AUTHOR CONTRIBUTIONS

J.-M.A.-C. performed analytic work, Voronoi model simulations and analyzed data. L.N.C. coordinated the research, performed the multiphase field simulations and analyzed data. J.E. performed the experiments and analyzed data. D.K. performed analytic work and analyzed data. L.G. devised and coordinated the research. All authors wrote the paper. J.-M.A.-C., L.C.N., J.E. and D.K. contributed equally to this work.

## AUTHOR INFORMATION

The authors declare no competing financial interests.

## METHODS

### Cell culture

Parental Madin-Darby Canine Kidney (MDCK) GII cells stably expressing E-cadherin-GFP [35] (kindly provided by M. Gloerich, UMC Utrecht) were cultured in a 1 : 1 ratio of low glucose DMEM (D6046; Sigma-Aldrich, St. Louis, MO) and Nutrient Mixture F-12 Ham (N4888; Sigma-Aldrich, St. Louis, MO) supplemented with 10% fetal calf serum (Thermo Fisher Scientific, Waltham, MA), and 100 mg*/*mL penicillin/streptomycin, 37 ^°^C, 5% CO_2_. For experiments, cells were seeded on uncoated cover glasses, grew to confluence, and nuclei were live-stained with 2 *µ*g/mL Hoechst 34580 (Thermo Fisher, H21486) before imaging.

### Microscopy

Samples were imaged at high resolution on a home-build optical microscope setup based on an inverted Axiovert200 microscope body (Zeiss), a spinning disk unit (CSU-X1, Yokogawa), and an emCCD camera (iXon 897, Andor). IQ-software (Andor) was used for setup-control and data acquisition. Illumination was performed using fiber-coupling of different lasers [405 nm (CrystalLaser) and 488 nm (Coherent)]. Cells on over glasses were inspected with an EC Plan-NEOFLUAR 40 × 1.3 Oil immersion objective (Zeiss). Images were taken in three focal-planes within a distance of 352 nm for a maximal intensity projection.

### Analysis

#### Shape order parameter

Cell boundaries of confluent monolayers were analyzed using a maximum intensity projection of *z*−stack images. Cell segmentation and vertex analysis were performed using home-build Matlab scripts (Mathworks, Matlab R2018a). The number of nearest neighbors corresponds to the number of vertices surrounding a cell. The centroid of the polygon was calculated by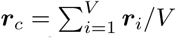, where *V* is the number of vertices and ***r***_*i*_ their positions. For each cell, the shape order was derived by using Eq.(2). On average, we analyzed 140 ± 31 cells per image. For the probability distribution of the shape order for each analyzed image, we choose a binning of 20 ranging from 0 to 1.

#### Coarse graining

The radius used to construct the coarse grained field given by Eq. (3), was chosen according to the typical cell radius *R*_cell_ = 7.4 ±1.9 *µ*m, calcualted as half of the average cell-cell nearest neighbor distance. For calculating the crossover point, we set the center point of the disk equal to the center point of the image. The radius of the disk in which the complex order parameters were averaged ranged from *R*_cell_ to half of the image size (176 × 176*µ*m^2^). For computing the nematic and hexatic coarse grained director field, we set the grid-distance to *R*_cell_.

#### Topological defects

Topological defects were identified by first interpolating the *p−*fold orientation field on a square 22 × 22 grid by means of the coarse-graining procedure in Eq. (3) and then computing the winding number along each unit cell. That is:

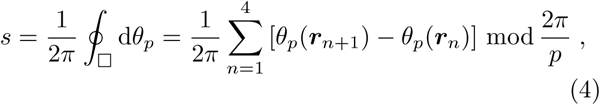

where the symbol □ denotes a square unit cell in the interpolation grid and the mod operator constraints the difference *θ*_*p*_(***r***_*n*+1_)−*θ*_*p*_(***r***_*n*_) in the interval [−2*π/p*, 2*π/p*].

### Statistics

In total, 68 images of confluent monolayers (nine coverslips, three independent experiments) were taken and analyzed. In total, 9496 cells were considered for the analysis.

### Numerical simulations

We make use of two different numerical models for ET previously intrduced in literature: *(i)* the multiphase field model and *(ii)* the Voronoi model.

#### Multiphase field model

This model has been used to study the dynamics of confluent cell monolayers [17] and the mechanics of cell extrusion [18]. It is a continuous model where each cell is described by a concentration field *φ*_*c*_ = *φ*_*c*_(***r***), with *c* = 1, 2 … *N*_cell_ and *N*_cell_ the total number of cells. The equilibrium state is defined by the free energy ℱ = ∫d*A f* where the free energy density *f* is given by

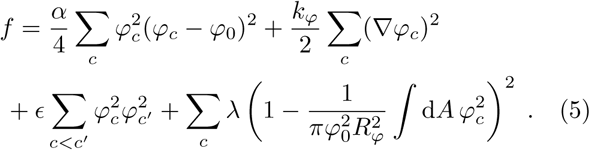

Here *α* and *k*_*φ*_ are material parameters which can be used to tune the surface tension 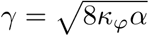 and the interfacial thickness 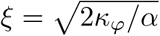 of isolated cells and thermodynamically favor spherical cell shapes. The constant *ϵ* captures the repulsion between cells. The concentration field is large (i.e. *φ*_*i*_ ≃*φ*_0_) inside the cells and zero outside. The contribution proportional to *λ* in the free energy enforces cell incompressibility whose nominal radius is given by *R*_*φ*_. The phase field *φ*_*i*_ evolves according to the Allen-Cahn equation

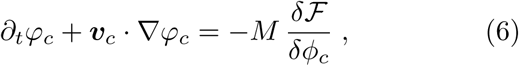

where ***v***_*i*_ = *v*_0_(cos *θ*_*c*_ ***e***_*x*_+sin *θ*_*c*_ ***e***_*y*_) is the velocity at which the *c −*th cell self-propels, with *v*_0_ a constant speed and *θ*_*c*_ an angle. The latter evolves according to the stochastic equation

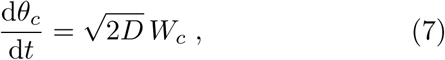

where *D* is a constant controlling noise diffusivity and *W*_*c*_ = *W*_*c*_(*t*) is a Wiener process. The constant *M* in Eq. (6) is the mobility measuring the relevance of thermodynamic relaxation with respect to non-equlibrium cell migration. Eq. (6) is solved with a finite-difference approach through a predictor-corrector finite difference Euler scheme implementing second order stencil for space derivatives [19]. Simulation details and scaling to physical units are given in Table I.

**Table I.** Physical scaling of simulation parameters. The table provides the parameters used to perform simulations for both the multiphase field and the Voronoi model, together with their dimensions and scaling to physical units. For the multiphase field model, scaling is performed by equating the mean cell radius *R*_cell_ (≃ 7.4*μ*m) measured in experiments with the nominal cell radius *R* and a typical migration speed of cells in MDCK monolayers [6] (≃ 2*μ*m h^*−*1^) with that measured in our simulations (≃ 0.0011Δ*x/*Δ*t*). This allows us to find the physical scaling of the lattice grid unit Δ*x* and the iteration unit Δ*t*. For the Voronoi model, we equated the mean cell radius *R*_cell_ in experiments with that measured in simulations (≃ 1). The time-step was derived with the same procedure as described for the multiphase field model.In the table, simulation values are given in both lattice and physical units, in column four and five, respectively. Notice that we did not introduce an energy scale as this cancels out with the mobility parameter *M* in Eq. (6) and *μ* in Eq. (8), respectively.

| Numerical model |  |  |  |  |
| --- | --- | --- | --- | --- |
| *Multiphase field model* |  |  |  |  |
| Model parameter |  | Dimension | Simulation value(s) | Physical scaling |
| *Lattice parameters* | $N_{\text{cell}}$ | — | 361 | — |
| | $\Delta x$ | $L$ | 1 | $0.685 \mu\text{m}$ |
| | $\Delta t$ | $T$ | 1 | $1.414 \text{ s}$ |
| | $L_x, L_y$ | $L$ | 380 | $246.6 \mu\text{m}$ |
| *Free energy parameters* | $M\alpha$ | $1/T$ | 0.006 | $0.0042 \text{ s}^{-1}$ |
| | $Mk_\phi$ | $L^2/T$ | 0.012 | $0.0040 \mu\text{m}^2 \text{ s}^{-1}$ |
| | $M\epsilon$ | $1/T$ | 0.01 | $0.0071 \text{ s}^{-1}$ |
| | $M\lambda$ | $1/T$ | 600 | $424, 4 \text{ s}^{-1}$ |
| | $\varphi_0$ | — | 2.0 | — |
| | $R$ | $L$ | 10.86 | $7.4 \mu\text{m}$ |
| | $\xi = \sqrt{2k_\phi/\alpha}$ | $L$ | 2 | $1.37 \mu\text{m}$ |
| | $M\gamma = M\sqrt{8/9k_\phi\alpha}$ | $L/T$ | 0.008 | $0.0039 \mu\text{m s}^{-1}$ |
| *Dynamical equation parameter* | $D_r^{pf}$ | $1/T$ | 0.0001 | $0.00007 \text{ s}^{-1}$ |
| | $v_0$ | $L/T$ | 0.0035 | $0.00169 \mu\text{m s}^{-1}$ |
| *Dimensionless numbers* | Peclet number $Pe = v_0/(D_r R)$ | — | 3.22 | — |
| | Cell deformability $d = \epsilon/\alpha$ | — | 1.66 | — |
| *Voronoi model* |  |  |  |  |
| *Lattice parameters* | $N_{\text{cell}}$ | — | 22500 | — |
| | $\Delta t$ | $T$ | 0.01 | $0.53 \text{ s}$ |
| | $L_x, L_y$ | $L$ | 150 | $2220 \mu\text{m}$ |
| *Energy parameters* | $\mu K_A$ | $1/(L^2 T)$ | 1 | $0.0086 \mu\text{s}^{-1}$ |
| | $\mu K_P$ | $1/T$ | 1 | $0.98 \text{ s}^{-1}$ |
| | $A_0$ | $L^2$ | 1 | $219.04 \mu\text{m}^{-2}$ |
| | $P_0$ | $L$ | 3.9 | $57.72 \mu\text{m}$ |
| *Dynamical equation parameter* | $v_0$ | $L/T$ | 0.1 | $27.8 \mu\text{m s}^{-1}$ |
| | $D'_r$ | $1/T$ | 1 | $0.019 \text{ s}^{-1}$ |
| *Dimensionless numbers* | Peclet $Pe = v_0/(D'_r \sqrt{A_0})$ | — | 0.5 | — |
| | Shape index $p_0 = P_0/\sqrt{A_0}$ | — | 3.95 | — |

#### Voronoi model

This model portrays a confluent tissue as a Voronoi tesselation of the plane [20]. Each cell is characterized by two dynamical variables: the position ***r***_*c*_ and the velocity ***v***_*c*_ = *v*_0_(cos *θ*_*c*_ ***e***_*x*_ + sin *θ*_*c*_ ***e***_*y*_) with *v*_0_ a constant speed and *θ*_*c*_ an angle, with *c* = 1, 2 … *N*_cell_ and *N*_cell_ the total number of cells. The dynamics of these variables is governed by the following set of ordinary differential equations

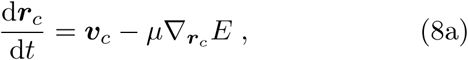

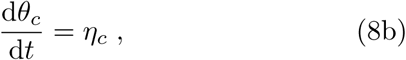

where *µ* is a mobility coefficient and *E* = *E*(***r***_1_, ***r***_2_ … ***r***_*N*_) is and energy function defined as

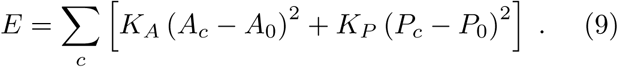

Here *A*_*c*_ and *P*_*c*_ are respectively the area and perimeter of each cell and *A*_0_ and *P*_0_ its preferred values. The variable *η*_*c*_ in Eq. (8b) is white noise, having zero mean and correlation function

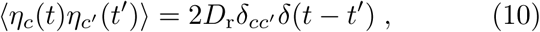

with *D*_r_ a rotational diffusion coefficient. Simulation details and scaling to physical units are given in Table I.

## Supplementary information

### S1. THE *p−*FOLD SHAPE TENSOR

#### A. Definition and basic properties

In this supplementary Section we explicit the relation between the *p*−fold shape tensor ***G***_*p*_, Eq. (1), and the complex order parameter *ψ*_*p*_, Eq. (2). To build up intuition, we start from observing that the standard rank−2 shape tensor for a *V* −sided polygon, is given by^8, 9^

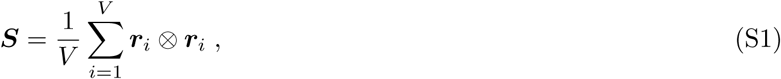

where, as in the main text, ***r***_*i*_ represents the coordinate of the *i*−th vertex with respect to the center of mass of the cell. The spectral theorem allows one to represent ***S***, as well as any other symmetric tensor, in terms of two irreducible components, one diagonal and the other traceless:

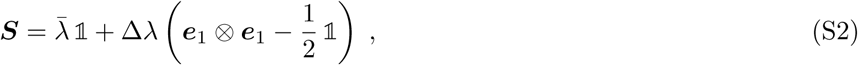

where we have set

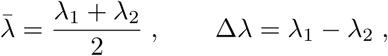

with *λ*_1_ *> λ*_2_ the two eigenvalues of ***S, e***_1_ = cos *ϑ* ***e***_*x*_ + sin *ϑ* ***e***_*y*_ the unit eigenvector associated with the largest eigenvalue *λ*_1_ and 𝕝 the rank–2 identity tensor. The two terms in Eq. (S2) entail information about the polygon’s size and anisotropy. The latter property can be further highlighted by introducing the tensor

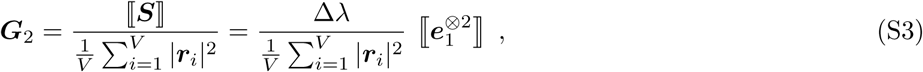

where the operator ⟦· · ·⟧ has the effect of rendering its argument traceless and symmetric^13, 14^ and the (· · ·)^⊗*p*^ implies a *p*−fold tensorial product of the argument with itself: i.e.

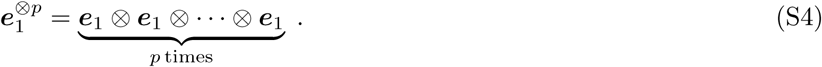

In two dimensions, the tensor ***G***_2_ has only two linearly independent components and expressing it in the basis {***e***_*x*_, ***e***_*y*_} readily gives

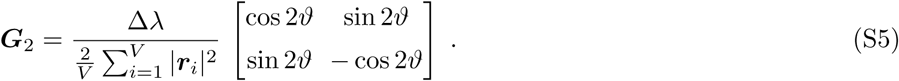

Furthermore, explicitly diagonalizing Eq. (S1) gives

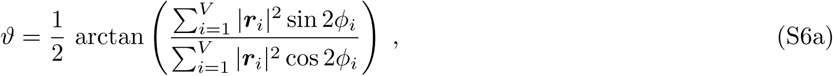

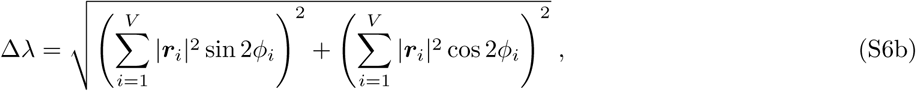

where *φ*_*i*_ = arctan(*y*_*i*_*/x*_*i*_) denotes the angular position of the *i*−th vertex with respect of the centre of mass (see Fig. 1e of the main text). This construction implies that all components of the tensor ***G***_2_ are proportional to either the real or imaginary part of the complex order parameter

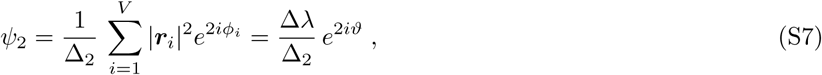

so that

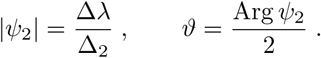

Now, the same construction can be carried out for a generic rank−*p* shape tensor, by defining

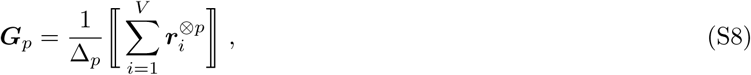

Where 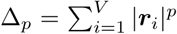. As for the rank−2 tensor defined in Eq. (S3), this tensor has only two linearly independent components, that are

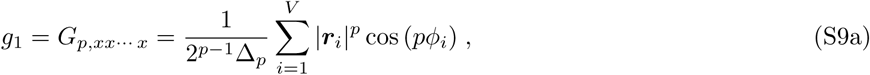

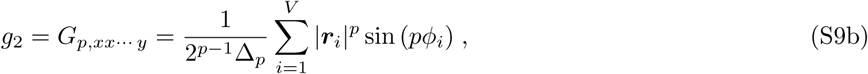

and can be cast as in Eq. (S3), that is

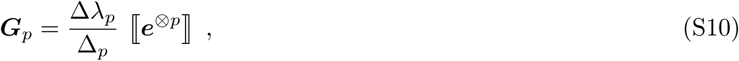

where the positive scalar Δ*λ*_*p*_ and the unit vector ***e*** = cos *ϑ*_*p*_ ***e***_*x*_ + sin *ϑ*_*p*_ ***e***_*y*_ are analogous to the difference *λ*_1_ −*λ*_2_, quantifying the anisotropy of the polygon, and the eigenvector ***e***_1_ associated with the largest eigenvalue. This problem ultimately relies on a generalization of the spectral theorem for tensors whose rank is larger than two. A possible strategy to achieve such as generalization was proposed by Virga in the context of rank −3 tensors^15^ and consists of defining *ϑ*_*p*_ as the inclination of a *p*−legged star oriented in such a way to maximize the probability of finding a vertex of the polygon in the direction of either one of the legs. The latter task is equivalent to solving the system of equations

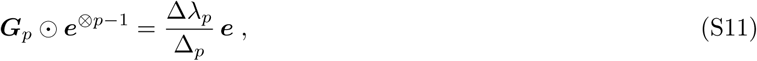

where ⊙ denotes a contraction of all matching indices of the two tensors on the left hand side. After some lengthy calculations, partially summarized in Sec. S1 B, one finds

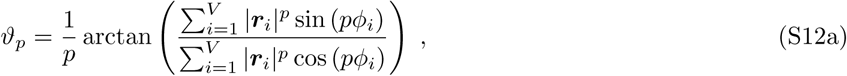

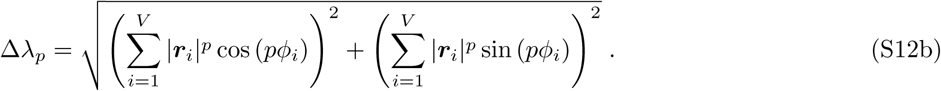

As in the case of the rank−2 shape tensor, one can then express all components of ***G***_*p*_ in terms of the real and imaginary parts of the *p*−fold complex order parameter

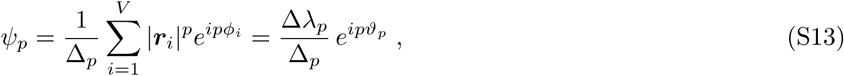

so that

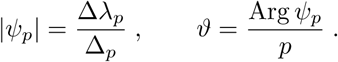

#### B. Derivation of Eqs. (S12)

For sake of completeness, here we elaborate on the solution of Eq. (S11), leading to Eqs. (S12). The strategy, pioneered in Ref.^15^, consists of mapping the diagonalization of a rank−*p* tensor to an optimization problem where Δ*λ*_*p*_ ∈ ℝ is the Lagrange multiplier subjected to the constraint 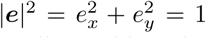. This task requires computing the tensorial power ***e***^⊗*p*−1^, which, in turn, amounts to constructing all possible order −(*p* −1) products of *e*_*x*_ and *e*_*y*_. The latter is facilitated by the fact that, as previously stated, the two-dimensional tensor ***G***_*p*_ has only two linearly independent components, proportional to the functions *g*_1_ and *g*_2_ introduced in Eqs. (S9). In particular, depending on whether the number of *y*−indices of the generic element 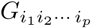, with *i*_*p*_ = {*x, y*}, is even or odd, the element is proportional to *g*_1_ and *g*_2_ respectively. Taken together, the aforementioned considerations result into the following expressions for the components of the ***e*** vector:

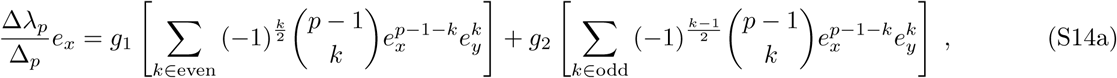

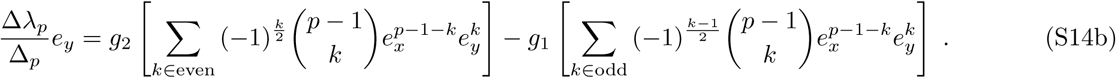

Despite their apparently complexity, these equations can be considerably simplified leading to

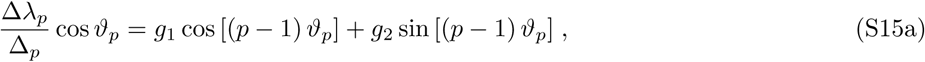

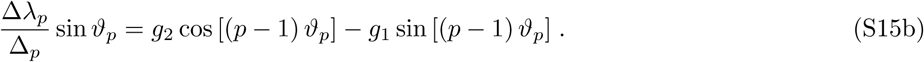

If *g*_1_ = 0, Eqs. (S15) reduces to

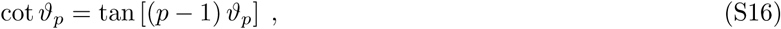

which has 2 *p* solutions in the range 0 ≤ *ϑ*_*p*_ *<* 2*π* given by

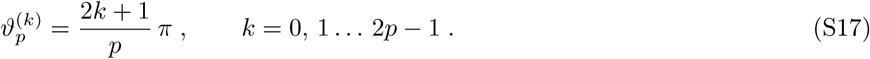

Conversely, when *g*_1_ ≠ 0, setting *ϱ* = *g*_2_*/g*_1_ and solving Eqs. (S15) with respect to *ϑ*_*p*_ gives

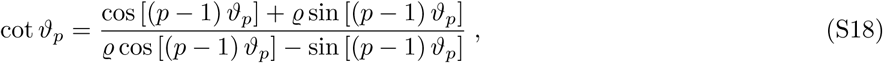

from which one can readily find

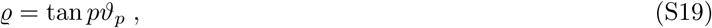

whose solution is given by

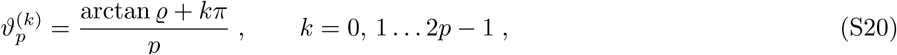

thereby completing the derivation of Eq. (S12a). To compute Δ*λ*_*p*_ one can use again Eqs. (S15) and express *g*_1_ and *g*_2_ in terms of coordinates. This gives, after some direct calculations

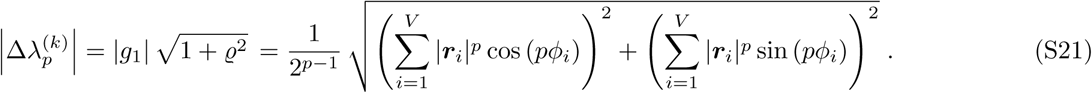

Note that, because of the periodicity of 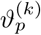, then 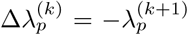, whereas the sign of 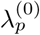 depends on *ϱ* and *g*_1_. Finally, to cast the tensor ***G***_*p*_ in the form given in Eq. (S10), one can write 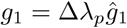 and 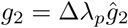 where

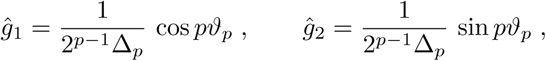

are the two independent components of ⟦ ***e***^⊗*p*^*⟧/ Δ*_*p*_. Then, using the expression of *ϑ*_*p*_ given in Eqs. (S17) and (S20), one obtains

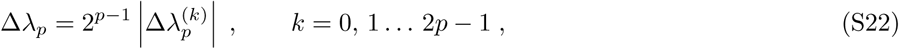

which completes the derivation of Eq. (S12b).

### S2. DEFECT REPRESENTATION IN *p*-ATIC LIQUID CRYSTALS

In two-dimensional liquid crystals topological defects consists of point-like singularities in the orientational field, that is points where the orientation of the director field is not univocally defined, and can be classified in terms of the winding number *s* defined in the main text. In liquid crystals with *p*−fold rotational symmetry, the latter is an integer multiple of the elementary winding number 1*/p*. By contrast, it is impossible to correctly describe a defect of winding number *s* = ±1*/p* in terms of an orientation field with rotational symmetry other than *p*−fold.

To substantiate this statement, we consider here the common case of a pair of ±1*/*2 disclinations in a nematic liquid crystal (*p* = 2), respectively located at positions ***r***_+_ = *x*_+_***e***_*x*_ + *y*_+_***e***_*y*_ and ***r***_−_ = *x*_−_***e***_*x*_ + *y*_−_***e***_*y*_. The far-field configuration the phase *ϑ*_2_ = Arg(Ψ_2_)*/*2 is given by

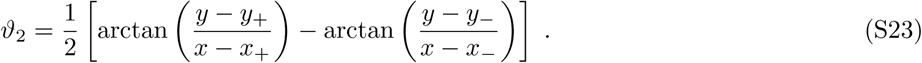

In turn, the 2−fold orientation field can be visualized as a the standard headless nematic director − i.e. a 2−legged star − as in Fig. 1a of the main text. Now, as illustrated in Fig. 3e, attempting to describe the same 2−fold symmetric configuration with a, say, 1−fold symmetric orientation filed − i.e. a standard vector field − results in a discontinuity of magnitude *π* of the associated phase *ϑ*_1_ across the *x*−axis.

The same issue occurs while attempting to describe a pair of *s* = ±*n/p* defects (with *n* a real number) in by means of a *q*−fold orientation filed, with *q < p*. In this case, the far-field configuration of the phase *ϑ*_*p*_ is given by

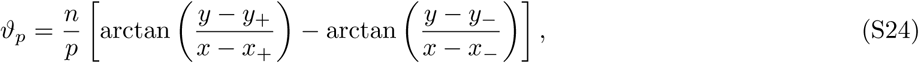

and it can be graphically represented by a *p*−legged star oriented at angles *ϑ*_*p*_ + 2*πn/p*, with *n* = 1, 2 … *p*, so that the order parameter Ψ_*p*_ = |Ψ_*p*_| exp *ipϑ*_*p*_ is continuous everywhere, but at the defect position. Next, we attempt to describe the same configuration in terms of the order parameter Ψ_*q*_ = Ψ_*q*_ exp *iqϑ*_*q*_ corresponding to *q*−legged stars oriented at an angle *ϑ*_*q*_ + 2*πn/q*, with *n* = 1, 2 … *q < p*. For the purpose of this discussion, and without loss of generality, we set *y*_+_ = *y*_−_ = *y*_0_ and we compute the variation of Ψ_*q*_ while crossing the line the axis *y* = *y*_0_ in the region comprised between the two defects (*x*_−_ *< x < x*_+_). Since the *q*−legged star associated with the order parameter Ψ_*q*_ is invariant under rotations by 2*π/q*, the inclination of the leg closer to the *x*−axis undergoes a discontinuity of magnitude

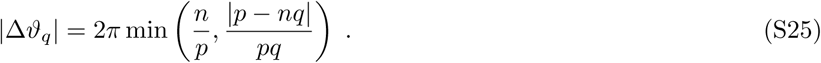

Thus, the field *ϑ*_*q*_ is continuous everywhere, but at the defect position (|Δ*ϑ*_*q*_| = 2*πm/p* with *m* any natural number) only when *p* = *q* or *nq* is an integer multiple of *p*. In particular, describing a defect of winding number *s* = 1*/*6 (*n* = 1 and *p* = 6) by means of a nematic field with *q* = 2 would result into a jump of magnitude |Δ*ϑ*_2_| = *π/*3 as shown in Fig. 3e in the main text. The resulting configuration of the nematic director features a singular line connecting defects of opposite charge.

## References

[1] P. Friedl and D. Gilmour, Nat. Rev. Mol. Cell Bio. 10, 445– (2009).

[2] C. De Pascalis and S. Etienne-Manneville, Mol. Biol. Cell 28, 1833 (2017).

[3] G. Duclos, C. Erlenkämper, J.-F. Joanny, and P. Silberzan, Nat. Phys. 13, 58 (2016).

[4] T. B. Saw, A. Doostmohammadi, V. Nier, L. Kocgozlu, S. Thampi, Y. Toyama, P. Marcq, C. T. Lim, J. M. Yeomans, and B. Ladoux, Nature 544, 212 (2017).

[5] K. Kawaguchi, R. Kageyama, and M. Sano, Nature 545, 327 (2017).

[6] L. Balasubramaniam, A. Doostmohammadi, T. Saw, G. Narayana, R. Mueller, T. Dang, M. Thomas, S. Gupta, S. Sonam, A. Yap, Y. Toyama, R.-M. Mége, J. Yeomans, and B. Ladoux, Nat. Mater 20, 1156 (2021).

[7] J. Bigun, Optimal orientation detection of linear symmetry (Linköping University Electronic Press, 1987).

[8] M. Aubouy, Y. Jiang, J. A. Glazier, and F. Graner, Granul. Matter 5, 67 (2003).

[9] M. Asipauskas, M. Aubouy, J. A. Glazier, F. Graner, and Y. Jiang, Granul. Matter 5, 71 (2003).

[10] P. M. Chaikin and T. C. Lubensky, Principles of condensed matter physics (Cambridge University Press, Cambridge, 1995).

[11] G. Friedel, Ann. Phys. 9, 273 (1922).

[12] For tensors whose rank is higher than two, the property of being traceless implies that contracting any two indices of the tensor yields zero.

[13] L. Giomi, J. Toner, and N. Sarkar, under review (2021), 2111.04720.

[14] L. Giomi, J. Toner, and N. Sarkar, under review (2021), 2106.11957.

[15] E. G. Virga, Eur. Phys. J. E 38, 63 (2015).

[16] “Supplementary information,”.

[17] B. Loewe, M. Chiang, D. Marenduzzo, and M. C. Marchetti, Phys. Rev. Lett. 125, 038003 (2020).

[18] S. Monfared, G. Ravichandran, J. E. Andrade, and A. Doostmohammadi, 2108.07657, under review (2021).

[19] L. N. Carenza, G. Gonnella, A. Lamura, G. Negro, and A. Tiribocchi, Eur. Phys. J. E 42, 81 (2019).

[20] D. Bi, X. Yang, M. C. Marchetti, and M. L. Manning, Phys. Rev. X 6, 021011 (2016).

[21] Y.-W. Li and M. P. Ciamarra, Phys. Rev. Materials 2, 045602 (2018).

[22] M. Durand and J. Heu, Phys. Rev. Lett. 123, 188001 (2019).

[23] A. Pasupalak, L. Yan-Wei, R. Ni, and M. Pica Ciamarra, Soft Matter 16, 3914 (2020).

[24] C. Kittel and P. McEuen, Introduction to solid state physics, Vol. 8 (Wiley New York, 1996).

[25] F. C. Keber, E. Loiseau, T. Sanchez, S. J. DeCamp, L. Giomi, M. J. Bowick, M. C. Marchetti, Z. Dogic, and A. R. Bausch, Science 345, 1135 (2014).

[26] S. J. Streichan, M. F. Lefebvre, N. Noll, E. F. Wieschaus, and B. I. Shraiman, Elife 7, e27454 (2018).

[27] P. Guillamat, C. Blanch-Mercader, K. Kruse, and A. Roux, bioRxiv (2020), 10.1101/2020.06.02.129262.

[28] Y. Maroudas-Sacks, L. Garion, L. Shani-Zerbib, A. Livshits, E. Braun, and K. Keren, Nature Physics 17, 251 (2021).

[29] J.-M. Armengol-Collado, L. N. Carenza, and L. Giomi, in preparation (2021).

[30] J. T. Blankenship, S. T. Backovic, J. Sanny, O. Weitz, and J. A. Zallen, Dev. Cell 11, 459 (2006).

[31] M. Rauzi, Phil. Trans. R. Soc. B 375, 20190552 (2020).

[32] A. Haeger, S. Alexander, M. Vullings, F. M. Kaiser, C. Veelken, U. Flucke, G. E. Koehl, M. Hirschberg, M. Flentje, R. M. Hoffman, et al., J. Exp. Med> 217 (2020), 10.1084/jem.20181184.

[33] X. Serra-Picamal, V. Conte, R. Vincent, E. Anon, D. T. Tambe, E. Bazellieres, J. P. Butler, J. J. Fredberg, and X. Trepat, Nat. Phys. 8, 628 (2012).

[34] L. A. Hoffmann, L. N. Carenza, J. Eckert, and L. Giomi, Sci. Adv. in press (2022), 2105.15200.

[35] S. Yamada, S. Pokutta, F. Drees, W. I. Weis, and W. J. Nelson, Cell 123, 889 (2005).

